# Centrifuge: rapid and sensitive classification of metagenomic sequences

**DOI:** 10.1101/054965

**Authors:** Daehwan Kim, Li Song, Florian P. Breitwieser, Steven L. Salzberg

## Abstract

Centrifuge is a novel microbial classification engine that enables rapid, accurate and sensitive labeling of reads and quantification of species on desktop computers. The system uses an indexing scheme based on the Burrows-Wheeler transform (BWT) and the Ferragina-Manzini (FM) index, optimized specifically for the metagenomic classification problem. Centrifuge requires a relatively small index (4.2 GB for 4,078 bacterial and 200 archaeal genomes) and classifies sequences at very high speed, allowing it to process the millions of reads from a typical high-throughput DNA sequencing run within a few minutes. Together these advances enable timely and accurate analysis of large metagenomics data sets on conventional desktop computers. Because of its space-optimized indexing schemes, Centrifuge also makes it possible to index the entire NCBI non-redundant nucleotide sequence database (a total of 109 billion bases) with an index size of 69 GB, in contrast to k-mer based indexing schemes, which require far more extensive space. Centrifuge is available as free, open-source software from www.ccb.jhu.edu/software/centrifuge

## Background

Microbes such as archaea and bacteria are found virtually everywhere on earth, from soils and oceans to hot springs and deep mines [1]. They are also abundant on and inside living creatures, including a variety of niches on the human body such as the skin and the intestinal tract [2]. These invisible life forms perform a vast range of biological functions; they are indispensable for the survival of many species; and they maintain the ecological balance of the planet. Many millions of prokaryotic species exist [3], although only a small fraction of them (<1% in soil and even fewer in the ocean) can be isolated and cultivated [4]. High-throughput sequencing of microbial communities, known as metagenomic sequencing, does not require cultivation and therefore has the potential to provide countless insights into the biological functions of microbial species and their effects on the visible world.

In 2004, the RefSeq database contained 179 complete prokaryotic genomes, a number that grew to 954 genomes by 2009 and to 4,278 by December 2015. Together with advances in sequencing throughput, this ever-increasing number of genomes presents a challenge for computational methods that compare DNA sequences to the full database of microbial genomes. Analysis of metagenomics samples, which contain millions of reads from complex mixtures of species, necessitates a compact and scalable indexing scheme for classifying these sequences quickly and accurately. Most of the current metagenomics classification programs either suffer from slow classification speed, a large index size, or both. For example, machine-learning based approaches such as the Naive Bayes Classifier (NBC) [5] and PhymmBL [6, 7] classify less than 100 reads per minute, which is too slow for datasets that contain millions of reads. In contrast, the pseudoalignment approach employed in Kraken [8] processes reads far more quickly, more than 1 million reads per minute, but its exact k-mer matching algorithm requires a large index. For example, Kraken's 31–mer database requires 93 GB of memory (RAM) for 4,278 prokaryotic genomes, considerably more memory than today's desktop computers contain.

Fortunately, modern read-mapping algorithms such as Bowtie [9, 10] and bwa [11, 12] have developed a data structure that provides very fast alignment with a relatively small memory footprint. We have adapted this data structure, which is based on the BurrowsWheeler transform [13] and the Ferragina-Manzini (FM) index [14], to create a metagenomics classifier, Centrifuge, that can efficiently store large numbers of genome sequences, taxonomical mappings of the sequences, and the taxonomical tree. Centrifuge is open-source software freely available to download at www.ccb.jhu.edu/software/centrifuge.

## Core algorithmic methods

### Database Sequence Compression

We implemented memory-efficient indexing schemes for the classification of microbial sequences based on the FM index, which also permits very fast search operations. We further reduced the size of the index by compressing genomic sequences and building a modified version of the FM index for those compressed genomes, as follows. First, we observed that for some bacterial species, large numbers of closely related strains and isolates have been sequenced, usually because they represent significant human pathogens. Such genomes include *Salmonella enterica* with 138 genomes, *Escherichia coli* with 131 genomes, and *Helicobacter pylori* with 73 genomes available (these figures represent the contents of RefSeq as of December 2015). As expected, the genomic sequences of strains within the same species are likely to be highly similar to one another. We leveraged this fact to remove such redundant genomic sequences, so that the storage size of our index can remain compact even as the number of sequenced isolates for these species increases.

Figure 1 illustrates how we compress multiple genomes of the same species by storing near-identical sequences only once. First, we choose the two genomes (G1 and G2 in the figure) that are most similar among all genomes. We define the two most similar genomes as those that share the greatest number of k-mers (using k=53 for this study) after k-mers are randomly sampled at a rate of 1% from the genomes of the same species. In order to facilitate this selection process, we used Jellyfish [15] to build a table indicating which k-mers belong to which genomes. Using the two most similar genomes allows for better compression as they tend to share larger chunks of genomic sequences than two randomly selected genomes. We then compared the two most similar genomes using nucmer [16], which outputs a list of the nearly or completely identical regions in both genomes. When combining the two genomes, we discard those sequences of G_2_ with ≥99% identity to G_1_, and retain the remaining sequences to use in our index. We then find the genome that is most similar to the combined sequences from G_1_ and G_2_, and combine this in the same manner as just described. This process is repeated for the rest of the genomes.

**Figure 1.**
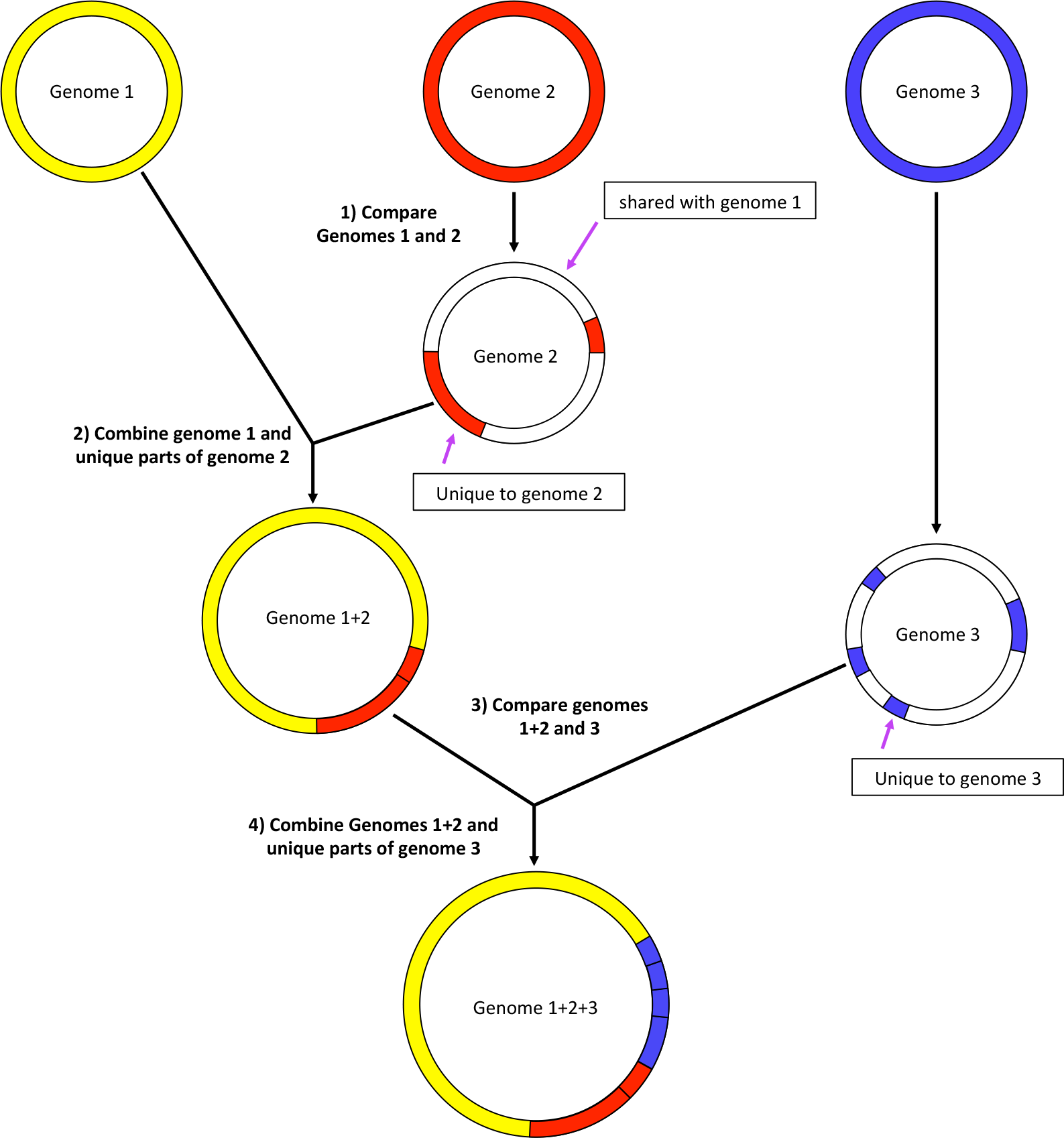
Compression of genome sequences before building the Centrifuge index.All genomes are compared and similarities are computed based on shared 53–mers. In the figure, genomes 1 and 2 are the most similar pair. Sequences in that are ≥99% identical between the two genomes are merged, and the remaining “unique” sequences from genome 2 are added to genome 1, created a merged (1+2) genome. Similarity between all genomes is recomputed using the merged genomes. Sequences <99% identical in genome 3 are then added to the merged genome, creating genome 1+2+3. This process repeats for the entire Centrifuge database until each merged genome has no sequences ≥99% identical to any other genome.

As a result of this concatenation procedure, we obtained dramatic space reductions for many species; e.g., the total sequence was reduced from 661 to 74 Mbp (11% of the original sequence size) in *S. enterica* and from 655 to 107 Mbp (16%) in *E. coli* (see Table 1). Overall, the number of base pairs from ~4,300 bacterial and archaeal genomes was reduced from 15 to 9.1 billion bases (Gb). The FM-index for these compressed sequences occupies 4.2 GB of memory, which is small enough to fit into the main memory (RAM) on a conventional desktop computer. As we demonstrate in the Supplementary Material, this compression operation has only a negligible impact on classification sensitivity and accuracy.

**Table 1.**
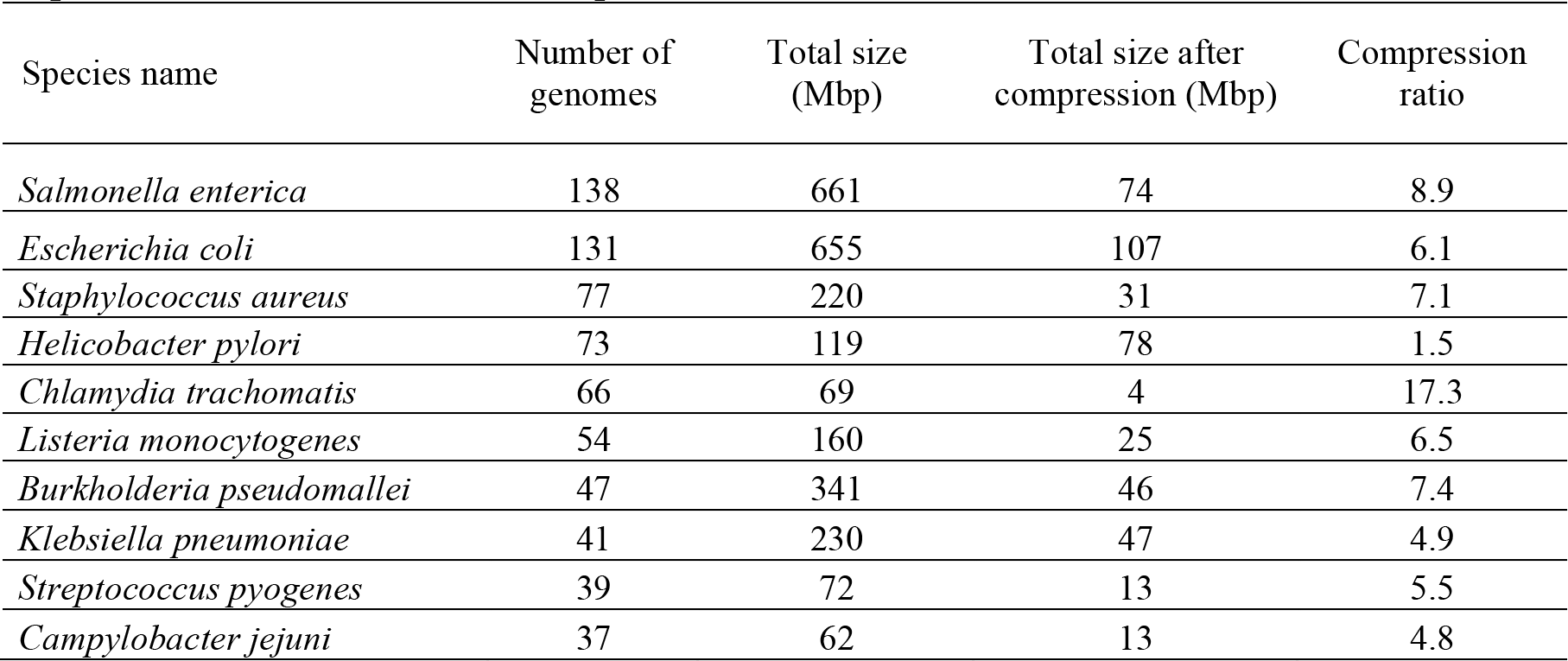
Compression ratios for 10 bacterial species that have multiple genomes fully sequenced and available in RefSeq.

### Classification based on the FM-index

The FM-index provides several advantages over k-mer based indexing schemes that store all k-mers in the target genomes. First, the size of the k-mer table is usually large; for example, Kraken’s k-mer table for storing all 31–mers in ~4,300 prokaryotic genomes occupies ~100 GB of disk space. Second, using a fixed value for k incurs a tradeoff between sensitivity and precision: classification based on exact matches of large k-mers (e.g., 31 bp) provides higher precision but at the expense of lower sensitivity, especially when the data being analyzed originates from divergent species. To achieve higher sensitivity, smaller k-mer matches (e.g., 20–25 bp) can be used, however this results in more false positive matches. The FM-index provides a means to exploit both large and small k-mer matches by enabling rapid search of k-mers of any length, at speeds comparable to those of k-mer table indexing algorithms (see Results).

Using this FM index, Centrifuge classifies DNA sequences as follows. Suppose we are given a 100–bp read (note the Centrifuge can just as easily process very long reads, assembled contigs from a draft genome, or even entire chromosomes). We search both the read (forward) and its reverse complement from right to left (3' to 5') as illustrated in Figure 2a. Centrifuge begins with a short exact match (16 bp minimum) and extends the match as far as possible. In the example shown in Figure 2a, the first 40 bp match exactly, with a mismatch at the 41^st^ base from the right. The rightmost 40 bp segment of the read is found in six species (A, B, C, D, E, F) that had been stored in the Centrifuge database. The algorithm then resumes the search beginning at the 42^nd^ base and stops at the next mismatch, which occurs at the 68^th^ base. The 26–bp segment in the middle of the read is found in species G and H. We then continue to search for mappings in the rest of the read, identifying a 32–bp segment that matches species G. Note that only exact matches are considered throughout this process, which is a key factor in the speed of the algorithm. We perform the same procedure for the reverse complement of the read, which in this example produces more mappings with smaller lengths (17, 16, 28, 18, and 17) compared to the forward strand.

**Figure 2.**
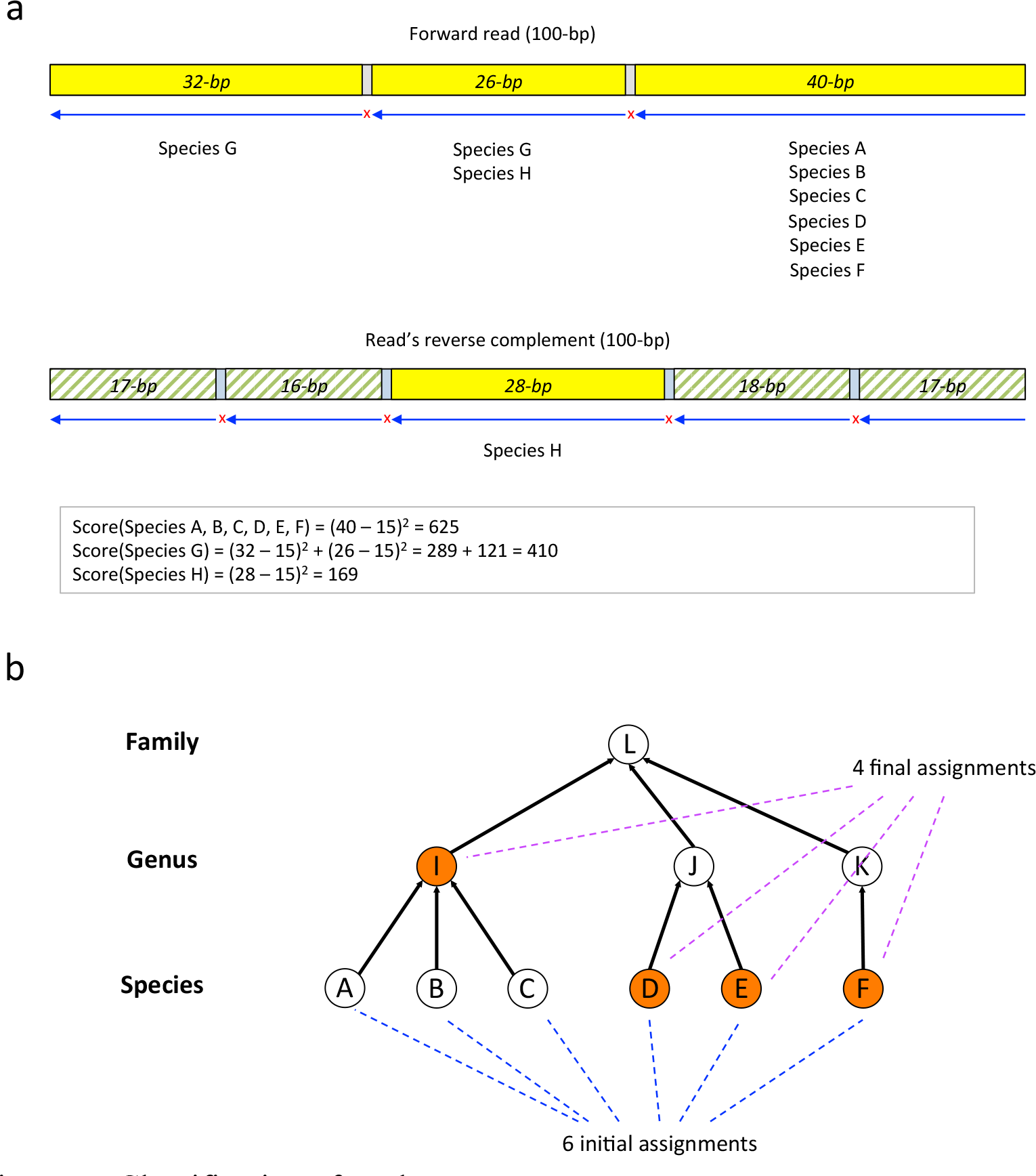
Classification of reads. The figure shows how the score for a candidate at the species level is calculated. Given a 100–bp read, both the read (forward) and its reverse complement from right to left are searched. Centrifuge first identifies a short exact match then continues until reaching a mismatch: the first 40 bp segment exactly matches six species (A, B, C, D, E, F), followed by a mismatch at the 41^st^ base, the second 26–bp segment matches two species (G and H), followed by a mismatch at the 68^th^ base, and the third 32-bp segment matches only species G. This procedure is repeated for the reverse complement of the read. Centrifuge assigns the highest score (625) to species A, B, C, D, E, and F. Centrifuge then traverses up the taxonomic tree to reduce the number of assignments, first by considering the genus that includes the largest number of species, genus I, which covers species A, B, and C, and then replacing these three species with the genus. This procedure results in reducing the number of assignments to four (genus I plus species D, E, and F).

Based on the exact matches found in the read and its reverse complement, Centrifuge then classifies the read using only those mappings with at least one 22–bp match. Figure 2a shows three segment mappings on the forward strand read and one on the read’s reverse complement that meet this length threshold. Centrifuge then scores each species using the following formula, which assigns greater weight to the longer segments: 
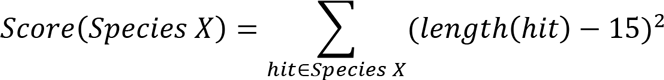
After assessing a variety of formulas, we empirically found that the sum of squared lengths of segments provides the best classification precision. For the example in Figure 2, species A, B, C, D, E, and F are assigned the highest score (625), based on the relatively long 40–bp exact match. Species G and H get lower scores because they have considerably shorter matches, even though each has 2 distinct matches. Note that H has mappings on both the read and its reverse complement, and in this case Centrifuge chooses the strand that gives the maximum score, rather than using the summed score on both strands,which might bias it towards palindromic sequences.

Centrifuge can assign a sequence to multiple taxonomic categories; by default, it allows up to five labels per sequence. (Note that this strategy differs from Kraken, which always chooses a single taxonomic category, using the lowest common ancestor of all matching species.) In Figure 2, six different species match the read equally well. In order to reduce the number of assignments, Centrifuge traverses up the taxonomic tree. First it considers the genus that includes the largest number of species, which in this example (Figure 2b) is genus I, which covers species A, B, and C. It then replaces these three species with the genus, thereby reducing the number of assignments to four (genus I plus species D, E, and F). If more than five taxonomic labels had remained, Centrifuge would repeat this process for other genuses and subsequently for higher taxonomic units until it reduced the number of labels to five or fewer.

The user can easily change the default threshold of five labels per sequence; for example, if this threshold is set to one, then Centrifuge will report only the lowest common ancestor as the taxonomic label, mimicking the behavior of Kraken. In the example shown in Figure 2, this label would be at the family level, which would lose some of the more specific information about which genuses and species the reads matched best.

If the size of the index is not a constraint, then the user can also use Centrifuge with uncompressed indexes, which classify reads using the same algorithm. Although considerably larger, the uncompressed indexes allow Centrifuge to classify reads at the strain or genome level; e.g., as *E. coli* K12 rather than just *E. coll.*

### Abundance analysis

In addition to per-read classification, Centrifuge performs abundance analysis at any taxonomic rank (e.g. strain, species, genus). Because many genomes share near-identical segments of DNA with other species, reads originating from those segments will be classified as multiple species. Simply counting the number of the reads that are uniquely classified as a given genome (ignoring those that match other genomes) will therefore give poor estimates of that species’ abundance. To address this problem, we define the following statistical model and use it to find maximum likelihood estimates of abundance through an Estimation-Maximization (EM) algorithm. Detailed EM solutions to the model have been previously described and implemented in the Cufflinks [17] and Sailfish [18] software packages.

Similar to how the Cufflinks calculates gene/transcript expressions, the likelihood for a specific configuration of species abundance α,given the read assignments *C,* is defined as follows: 
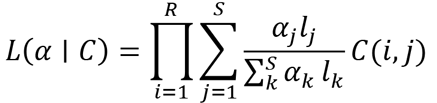
where *R* is the number of the reads, *S* is the number of species, α_*j*_ is the abundance of species *j,* summing up to 1 over all *S* species, *l_j_* is the average length of the genomes of species *j* and *C*(*i, j*) is 1 if read *i* is classified to species *j* and 0 otherwise.

To find the abundances α that maximize the likelihood function *L*(α), Centrifuge repeats the following EM procedure as also implemented in the Cufflinks until the difference between the previous estimate of abundances and the current estimate,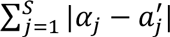, is less than 10^−10^.

Estimation (E-step):

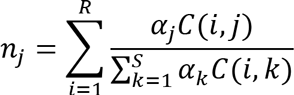
where *n_j_* is the estimated number of reads assigned to species *j*.

Maximization (M-step):

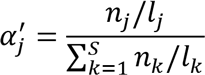
where 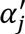is the updated estimate of species *j'*s abundance. α' is then used in the next iteration as α

## Results and discussion

We demonstrated the performance of Centrifuge in four different settings involving both real and simulated reads, and using several databases with different sizes, specifically one consisting of ~4,300 prokaryotic genomes (index name: *p*, index size: 4.2 GB), another with ~4,300 prokaryotic genomes plus human and viral genomes (*p+h+v*, 6.9 GB), and a third comprised of NCBI nucleotide sequences (nt, 69 GB). We compared the sensitivity and speed of Centrifuge to one of the leading classification programs, Kraken (v0.10.5– beta) [8]. We also included Megablast [19] in our assessment, as it is a very widely-used program that is often used for classification. In terms of both sensitivity and precision of classification, Centrifuge demonstrated similar accuracy to the other programs we tested. Centrifuge’s principal advantage is that it provides a combination of fast classification speed and low memory requirements, making it possible to perform large metagenomics analyses on a desktop computer using *p* or *p+h+v* index. For example, Centrifuge took only 47 minutes on a standard desktop computer to analyze 130 paired end RNA sequencing runs (a total of 26 GB) from patients infected with Ebola virus [20–22] as described below. Centrifuge’s efficient indexing scheme makes it possible to index the NCBI nucleotide collection (*nt*) database, which is a comprehensive set of sequences (> 36 million non-redundant sequences, ~110 billion bps) collected from viruses, archaea, bacteria, and eukaryotes, and enables rapid and accurate classification of metagenomics.

### Comparison of Centrifuge, Kraken, and Megablast on simulated reads from 4,278 prokaryotic genomes

We created a simulated read dataset from the 4,278 complete prokaryotic genomes in RefSeq [23] that were used to build the. From these genomes, we generated 10 million 100–bp reads with a per-base error rate of 3% using the Mason simulator, v0.1.2 [24].We used an error rate higher than found in Illumina reads (≤ 0.5%) in order to model the high mutation rates of prokaryotes. Reads were generated randomly from the entire data set; thus longer genomes had proportionally more reads. The full set of genomes is provided in Supplementary File 1. We built indexes for each of the respective programs. Kraken and Megablast require 100 GB and 25 GB of space (respectively) for their indexes. In contrast, Centrifuge requires only 4.2 GB to store and index the same genomes. The run-time memory footprint of MegaBlast is small (Table 2a) because it does not read the entire database into memory, in contrast to Kraken and Centrifuge. We classified the reads with Centrifuge, Kraken, and Megablast and calculated sensitivity and precision at the genus and species levels for each program (Table 2a). Centrifuge and Megablast often report multiple assignments for a given read, while Kraken instead reports the lowest common ancestor. To make our evaluation consistent across the programs, we only considered uniquely classified reads. Here we define sensitivity as the number of reads that are correctly classified divided by the total number of reads. Precision (also called positive predictive value) is defined as the number of correctly classified reads divided by the number of predictions made (i.e., reads that have no match and are not classified are not counted).

**Table 2.**
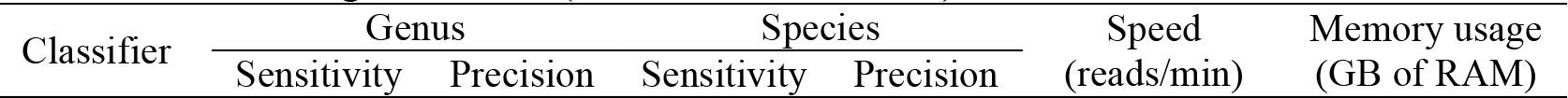
Classification sensitivity and precision for the Centrifuge, Kraken, and Megablast using simulated reads. In Centrifuge, we used only uniquely classified reads to compute accuracy. To measure speed, we used 10 million reads for Centrifuge and Kraken and 100,000 reads for Megablast. We ran all programs on a Linux system with 1 TB of RAM using one CPU (2.1 GHz Intel Xeon)

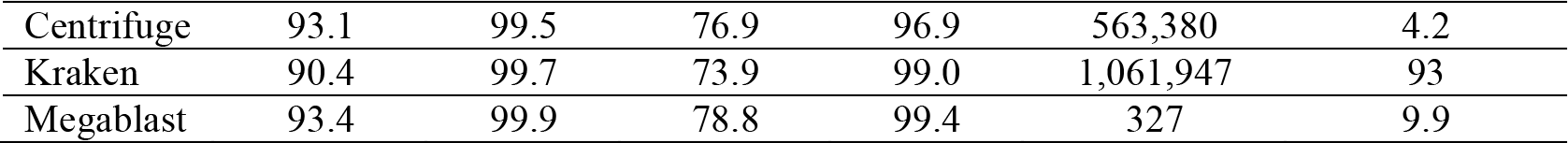

At the species level, Megablast provides the highest sensitivity at 78.8%, followed by Centrifuge (76.9%) and then Kraken (73.9%). Overall sensitivity is relatively low because many reads are assigned to multiple species and considered as unclassified in our evaluation. Megablast provides the highest precision, 99.4%, followed closely by Kraken at 99% then Centrifuge at 96.9%. At the genus level, Megablast provides the highest sensitivity at 93.4%, followed by Centrifuge (93.1%) and then by Kraken (90.4%). All three programs had near-perfect precision at the genus level, from 99.5–99.9%.

Kraken was the fastest program on this data, classifying about 1,062,000 reads per minute (rpm), followed by Centrifuge, which was approximately one-half as fast at 563,000 rpm. Megablast is far slower, processing only 327 rpm.

### Comparison of Centrifuge and Kraken performance on real datasets from sequencing reads of bacterial genomes

To test our method on real sequencing datasets, we downloaded 532 DNA sequencing datasets from the Sequence Read Archive (SRA). We selected them according to whether the SRA samples had been assigned a taxonomic identifier that belongs to a genus for which we have at least one genome in the database. All of the datasets were generated by whole-genome shotgun projects using recent Illumina platforms; 225 were sequenced on HiSeq and 307 on MiSeq instruments, with mean read lengths of 100 and 218 bps, respectively. Supplementary File 2 contains a complete list of the SRA identifiers, taxonomy IDs, number of reads and classification results. In total, this data contains over 560 million reads, with an average of 1,061,386 reads per sample (Supplementary Table 2).

**Figure 3.**
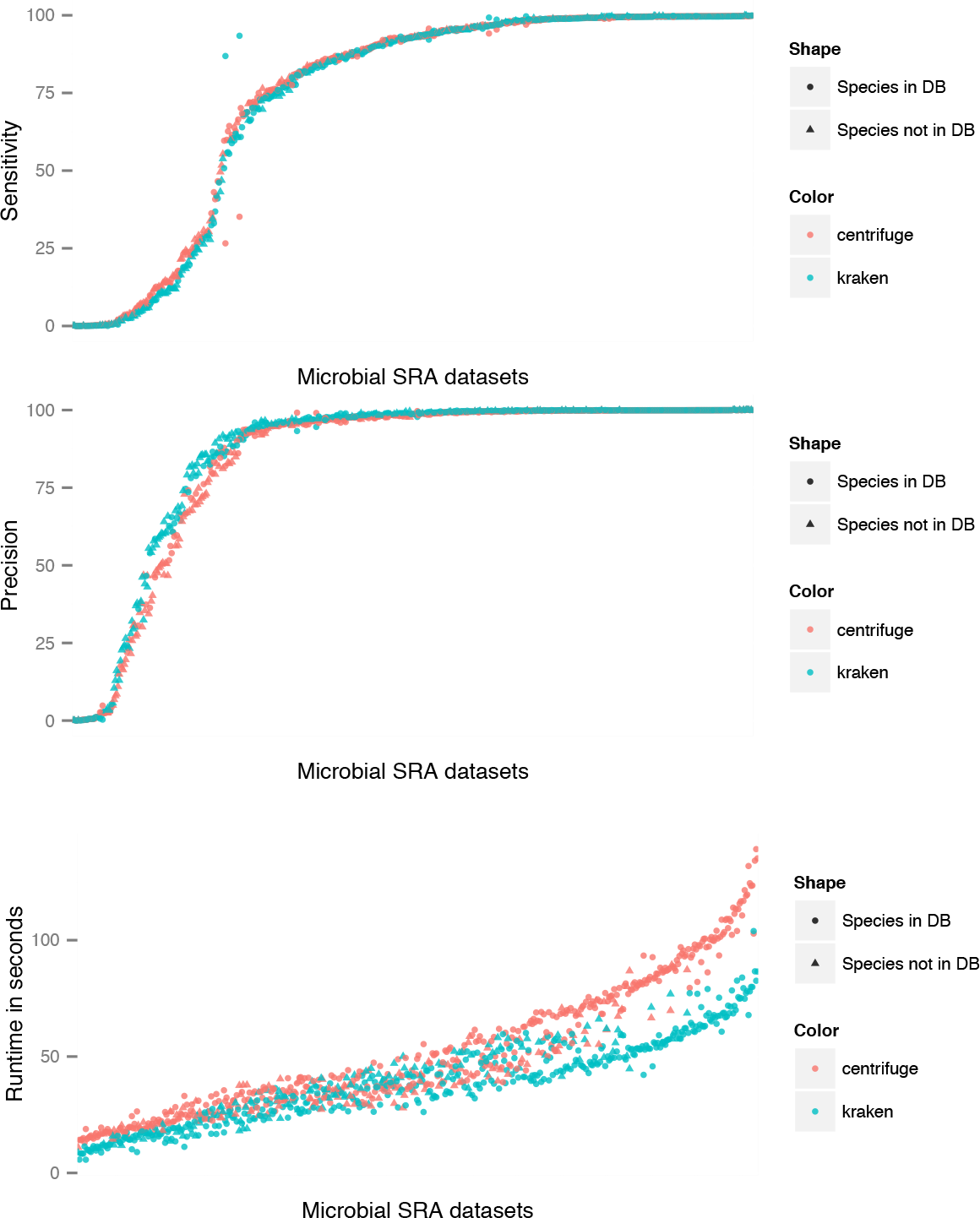
Results on 532 sequencing data sets from bacterial genomes retrieved from the Sequence Read Archive at NCBI. Each dot represents the results for one genome, with Centrifuge shown in orange and Kraken in teal. The uppermost plot shows sensitivity, computed as the percentage of reads classified as the correct species. The center plot shows precision, computed as the percentage of species-level classifications made by a program that were correct. The lower plot shows runtime measured in seconds.

For this experiment, we compared Centrifuge and Kraken but omitted Megablast because it would take far too long to run. Kraken was chosen as the standard for comparison because it demonstrated superior accuracy over multiple other programs in a recent comparison of metagenomic classifiers [25]. Figure 3 shows the results for both accuracy and speed using the database of ~4,300 prokaryotic genomes. On average, Centrifuge had slightly higher sensitivity (0.5% higher) than Kraken. Perhaps due to its use of a longer exact match requirement (31 bases), Kraken had slightly higher precision (1.5%) than Centrifuge. The lower accuracy of both programs on some data sets may be due to: 1) substantial differences between the genome that we have in the database and the strain that was sequenced, 2) numerous contaminating reads from the host or reagents, or 3) a high sequencing error rate for a particular sample.

Overall accuracy for both programs was very similar. Kraken was slightly faster, with an average run time (using 8 cores) of 39.4 seconds per genome, while Centrifuge required 50.9 seconds per genome.

### Application of Centrifuge for analyzing samples with Ebola virus and GB virus coinfections on a desktop

To demonstrate the speed, sensitivity, and applicability of Centrifuge on a real data set, we used data from the Ebola virus disease (EVD) outbreak. The 2013–2015 EVD outbreak in West Africa cost the lives of over 11,000 people as of August 26, 2015 (WHO Ebola Situation Report, apps.who.int/ebola/ebola-situation-reports.). In an international effort to research the disease and stop its spread, several groups sequenced the Ebola virus collected from patients’ blood samples and released their datasets online [20–22]. The genomic data was used to trace the disease and mutations in the Ebola genome, and inform further public health and research efforts. Lauck, et al. 2009 [26] reanalyzed one of the datasets (Gire, et al. 2009 [21]) in order to assess the prevalence and effect of GB virus C co-infection on the outcome of EVD.

We analyzed 130 paired-end sequencing runs from 49 patients reported in Gire, et al. 2009 [21] using Centrifuge to look for further co-infections. This dataset has a total of 97,097,119 reads (26 GB of FASTA files). The accession IDs of this data are provided in Supplementary File 3. For this analysis, we used the database containing all prokaryotic genomes (compressed), all viral genomes, and the human genome (total index size: 6.9 GB). Running on a desktop computer (quad-core, Intel Core i5–4460@3.2GHz with 8GB RAM), Centrifuge completed the analysis of all samples in 47 minutes with four threads. RNA-sequencing [27] requires more steps than DNA-sequencing, including the reverse-transcription of RNA to DNA molecules, which introduces sequencing biases and artifacts. In order to handle these additional sources of errors and remove spurious detections, we filtered the results to include only reads that have a matching length of at least 60bp on the 2x100–bp reads.

Figure 4 shows our classification results for the 49 patients. Centrifuge detects between 3,853 and 6,781,684 Ebola virus reads per patient. As reported by Lauck, et al., we also detected co-infection of Ebola virus and GB virus C in many of the patients. Centrifuge identified at least one read from this virus in 27 of the 49 patients; nine patient samples had 50 or more reads. Eleven patients had between one and 10 reads matching the Hepatitis B virus, and in one sample over one thousand reads aligned uniquely to this virus. This Hepatitis B co-infection has not been reported previously, demonstrating the inherent advantage of using a metagenomics classification tool, which can also detect off-target species.

**Figure 4.**
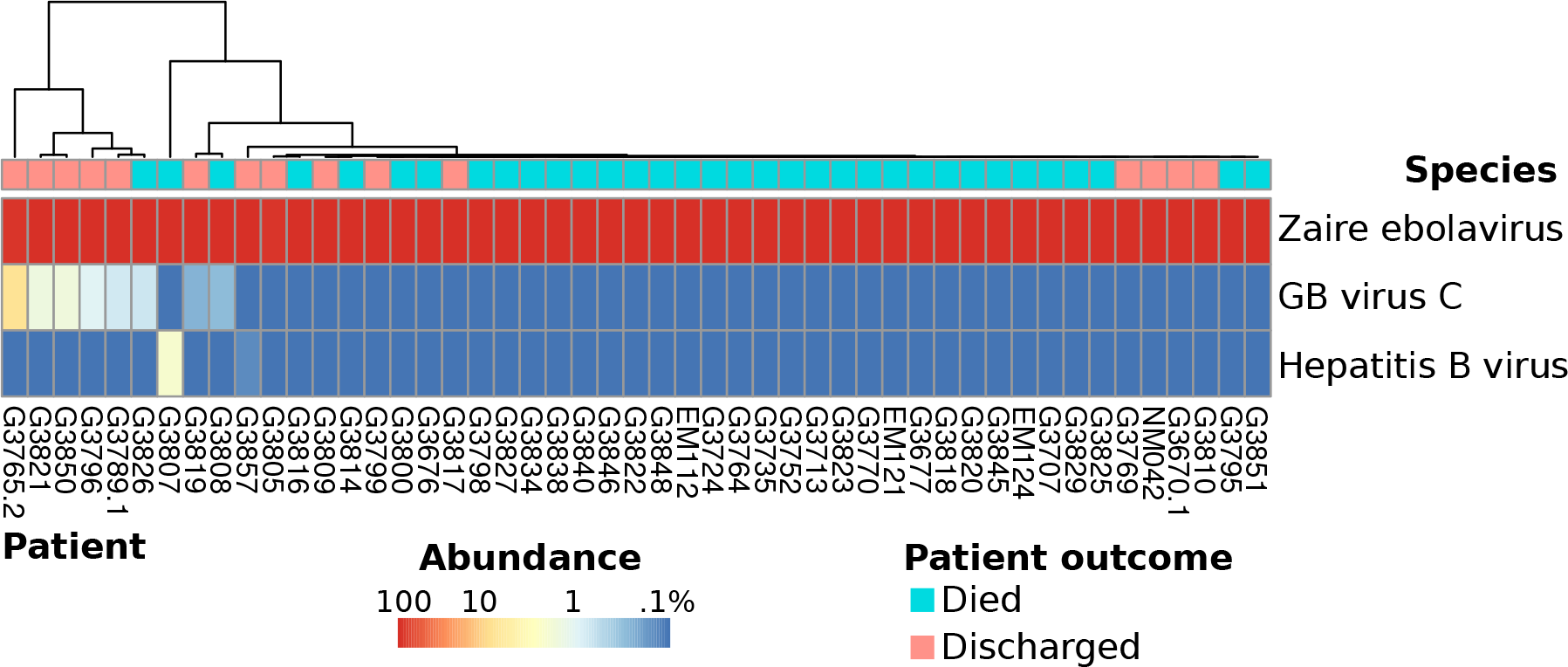
Heatmap of most abundant species in Ebola samples. The color scale encodes the abundance (number of unique reads normalized by genome size), from blue (less than .1% of normalized read count) to red (100% of reads). All species that have a normalized read count over 1% in any of the samples are shown. Zaire ebolavirus dominates the samples, however there is also a signal for other viruses in some of the patients - namely GB virus C and Hepatitis C virus.

### Centrifuge 's ability to index a comprehensive nucleotide sequence database

The NCBI nucleotide sequence database includes a comprehensive set of sequences collected from viruses, archaea, bacteria, and eukaryotes. Identical sequences have been removed to make it non-redundant, but even after this reduction the database contains over 36.5 million sequences with a total of ~109 billion bases (Gb). This rapidly growing database, called *nt*, enables the classification of sequencing data sets from hundreds of plant, animal, and fungal species as well as thousands of viruses and bacteria and many other eukaryotes. Metagenomics projects often include substantial quantities of eukaryotic DNA, and a prokaryotes-only index, as was used in the experiments above, cannot identify these species.

The challenge in using a much larger database is the far greater number of unique k-mers that must be indexed. For example, using Kraken's default k-mer length of 31 bp, the *nt* database contains approximately 57 billion distinct k-mers. Although it employs several elegant techniques to minimize space, Kraken still requires 12 bytes per k-mer, which means it would require an index size of 684 GB for the full *nt* database. Reducing the k-mer size helps only slightly: with k=22, Kraken would require an index of 520 GB.

Either of these indexes would require a specialized computer with very large main memory.

Centrifuge’s index is based on the space-efficient Burrows-Wheeler transform, and as a result it requires only 69 GB for the *nt* database, less than the raw sequence itself. Blast and Megablast are currently the only alternative methods that can classify sequences against the entire *nt* database; thus we compared Centrifuge with Megablast using our simulated read data, described above. Megablast uses a larger index, requiring 155 GB on disk, but it does not load the entire index into memory, and requires only 16 GB of RAM, while Centrifuge requires 69 GB. However, Centrifuge classified reads at a far higher speed: in our experiments on the Mason simulation data it processed ~372,000 reads/minute, over 3500 times faster than Megablast, which processed only 105 reads/minute. The classification precision and sensitivity of both programs were very high: Centrifuge’s precision and sensitivity were 99.5% and 89.9% respectively, while Megablast’s were 99.8% and 93.9%.

As a test of Centrifuge’s *nt* database, we used it to analyze sequences from a mixture of common fruits and vegetables sequenced using long-read single molecule technology.The mixture included more than a dozen common foods: grape, blueberry, yam (sweet potato), asparagus, cranberry, lemon, orange, iceberg lettuce, black pepper, wheat (flour), cherry tomato, pear, bread (wheat plus other ingredients), and coffee (beans). The “fruitshake” mixture was blended together, DNA was extracted, and sequencing was run on an Oxford Nanopore MinION. The number of reads generated from the fruitshake sample was 20,809, with lengths ranging from 90 to 13,174 bp and a mean length of 893 bps (Supplementary File 4 provides the reads in fastq format). Although MinION platforms produce much longer reads than Illumina platforms, MinIONs' high sequencing error rates (estimated at 15% [28]) prevent reads from containing long exact matches and increase the chance of noisy and incorrect matches. We initially labeled 8,236 reads using Centrifuge. In order to reduce false positive assignments for these error-prone reads, we filtered out those reads that scored ≤300 and had match lengths ≤50-bp, resulting in 3,617 reads ultimately classified. (Table 3 shows 14 species to which at least 5 reads are uniquely assigned, encompassing many of the species included in the sample such as wheat, tomato, lettuce, grape, barley, and pear. Note that as with any real sample, the true composition of the reads is unknown; we present these results here to illustrate (a) the use of the large *nt* database and (b) the use of Centrifuge on long, high-error-rate reads. Although apple was not known to be present in the sample, the 5 reads assigned to apple might have been due to similarity between the apple genome and the pear genome. 26 reads were identified as sheep and 8 as cow, which were confirmed separately by Blast searches. These could represent sample contamination or possibly contaminants in the sheep and cow assemblies. Missing species can be explained either by low abundance in the sample or because their genomes are substantially different from those in the Centrifuge *nt* database.

**Table 3.**
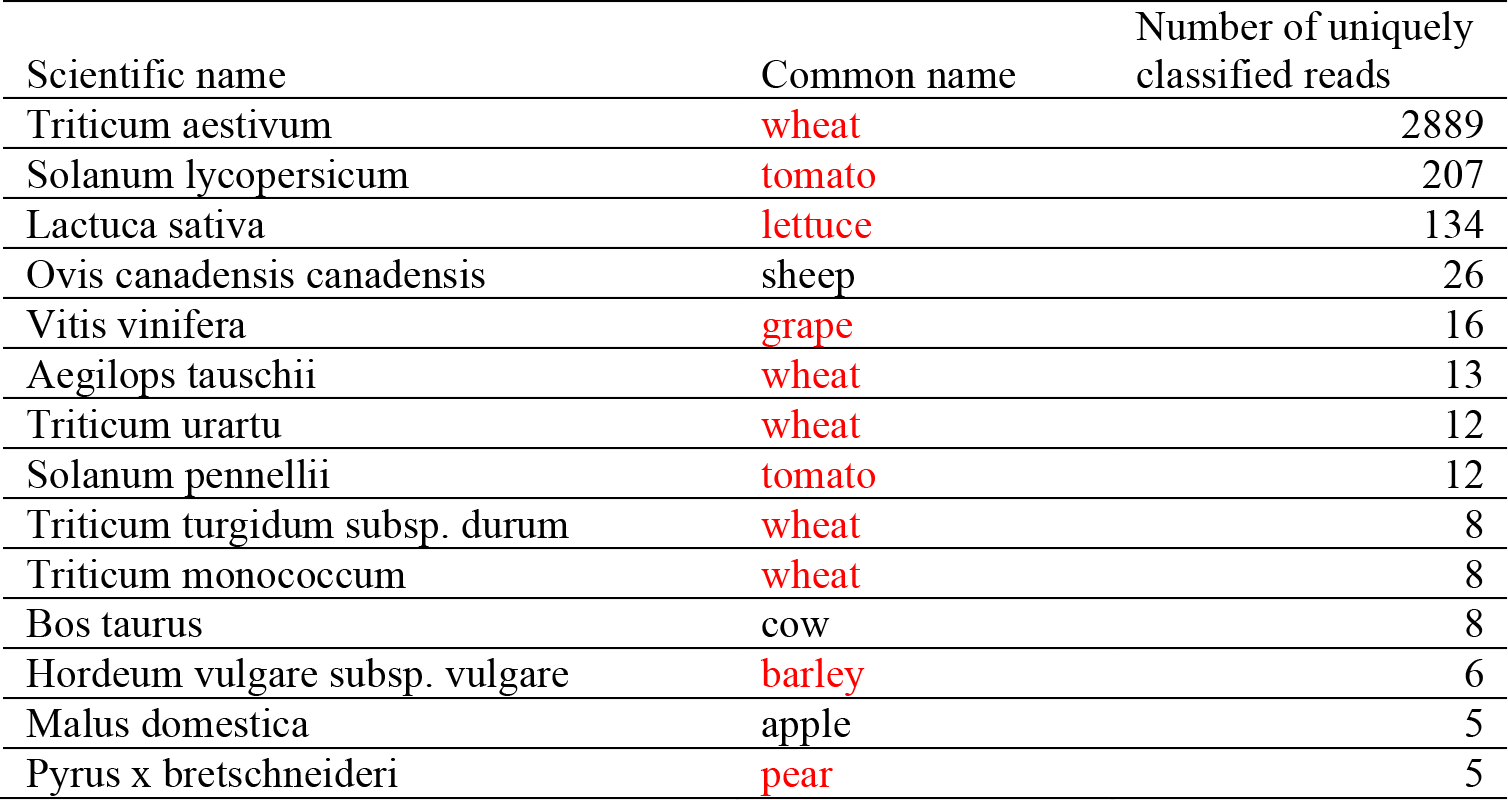
Classification of the fruitshake sample using Centrifuge’s nt database. The table shows 14 genomes to which at least 5 reads sequenced from the fruitshakesample were uniquely assigned. Common names in red represent species known to be present in the mixture.

| Scientific name | Common name | Number of uniquely classified reads |
| --- | --- | --- |
| *Triticum aestivum* | **wheat** | 2889 |
| *Solanum lycopersicum* | **tomato** | 207 |
| *Lactuca sativa* | **lettuce** | 134 |
| *Ovis canadensis canadensis* | sheep | 26 |
| *Vitis vinifera* | **grape** | 16 |
| *Aegilops tauschii* | **wheat** | 13 |
| *Triticum urartu* | **wheat** | 12 |
| *Solanum pennellii* | **tomato** | 12 |
| *Triticum turgidum* subsp. durum | **wheat** | 8 |
| *Triticum monococcum* | **wheat** | 8 |
| *Bos taurus* | cow | 8 |
| *Hordeum vulgare* subsp. vulgare | **barley** | 6 |
| *Malus domestica* | apple | 5 |
| *Pyrus x bretschneideri* | **pear** | 5 |

## Conclusions

Centrifuge is a rapid and sensitive classifier for microbial sequences with low memory requirements and a speed comparable to the fastest systems. Centrifuge classifies 10 million reads against a database of all complete prokaryotic and viral genomes within 20 minutes using one CPU core and requiring less than 8GB of RAM. Furthermore, Centrifuge can also build an index for NCBI’s entire *nt* database of non-redundant sequences from both prokaryotes and eukaryotes. The search requires a computer system with 128 GB of RAM, but runs over 3500 times faster than Megablast.

## Authors’ contributions

DK, LS, FPB, and SLS performed the analysis and discussed the results of Centrifuge. DK, LS, and FPB implemented Centrifuge. SLS collected and sequenced the fruitshake data. DK, FPB, LS, and SLS wrote the manuscript. All authors read and approved the final manuscript.

## Acknowledgements

This work was supported in part by NIH under grants R01–HG006677 and R01–GM083873, by the U. S. Army Research Office under grant W911NF–1410490, and by NSF under grant ABI–1356078.

## Additional files

Additional File 1: Supplementary material.

Additional File 2: Supplementary file 1.

Additional File 3: Supplementary file 2.

Additional File 4: Supplementary file 3.

## References

1. Keller M, Zengler K: Tapping into microbial diversity. Nature Reviews Microbiology 2004, 2:141–150.

2. Human Microbiome Project C: A framework for human microbiome research. Nature 2012, 486:215–221.

3. Schloss PD, Handelsman J: Status of the microbial census. Microbiol Mol Biol Rev 2004, 68:686–691.

4. Amann RI, Ludwig W, Schleifer KH: Phylogenetic identification and in situ detection of individual microbial cells without cultivation. Microbiol Rev 1995, 59:143–169.

5. Rosen G, Garbarine E, Caseiro D, Polikar R, Sokhansanj B: Metagenome fragment classification using N-mer frequency profiles. Adv Bioinformatics 2008, 2008:205969.

6. Brady A, Salzberg SL: Phymm and PhymmBL: metagenomic phylogenetic classification with interpolated Markov models. Nat Methods 2009, 6:673–676.

7. Brady A, Salzberg S: PhymmBL expanded: confidence scores, custom databases, parallelization and more. Nat Methods 2011, 8:367.

8. Wood DE, Salzberg SL: Kraken: ultrafast metagenomic sequence classification using exact alignments. Genome Biol 2014, 15:R46.

9. Langmead B, Trapnell C, Pop M, Salzberg SL: Ultrafast and memory-efficient alignment of short DNA sequences to the human genome. Genome Biol 2009, 10:R25.

10. Langmead B, Salzberg SL: Fast gapped-read alignment with Bowtie 2. Nature Methods 2012, 9:357–U354.

11. Li H, Durbin R: Fast and accurate short read alignment with BurrowsWheeler transform. Bioinformatics 2009, 25:1754–1760.

12. Li H, Durbin R: Fast and accurate long-read alignment with BurrowsWheeler transform. Bioinformatics 2010, 26:589–595.

13. Burrows M, Wheeler DJ: A Block-sorting Lossless Data Compression Algorithm. Technical Report 124 Palo Alto, CA: Digital Equipment Corporation 1994.

14. Ferragina P, Manzini G: Opportunistic data structures with applications.Foundations of Computer Science, 2000 Proceedings 41st Annual Symposium on 2000.

15. Marcais G, Kingsford C: A fast, lock-free approach for efficient parallel counting of occurrences of k-mers. Bioinformatics 2011, 27:764–770.

16. Kurtz S, Phillippy A, Delcher AL, Smoot M, Shumway M, Antonescu C, Salzberg SL: Versatile and open software for comparing large genomes. Genome Biol 2004, 5:R12.

17. Trapnell C, Williams BA, Pertea G, Mortazavi A, Kwan G, van Baren MJ, Salzberg SL, Wold BJ, Pachter L: Transcript assembly and quantification by RNA-Seq reveals unannotated transcripts and isoform switching during cell differentiation. Nat Biotechnol 2010, 28:511–515.

18. Patro R, Mount SM, Kingsford C: Sailfish enables alignment-free isoform quantification from RNA-seq reads using lightweight algorithms. Nat Biotechnol 2014, 32:462–464.

19. Zhang Z, Schwartz S, Wagner L, Miller W: A greedy algorithm for aligning DNA sequences. J Comput Biol 2000, 7:203–214.

20. Baize S, Pannetier D, Oestereich L, Rieger T, Koivogui L, Magassouba N, Soropogui B, Sow MS, Keita S, De Clerck H, et al:Emergence of Zaire Ebola virus disease in Guinea. N Engl J Med 2014, 371:1418–1425.

21. Gire SK, Goba A, Andersen KG, Sealfon RS, Park DJ, Kanneh L, Jalloh S, Momoh M, Fullah M, Dudas G, et al:Genomic surveillance elucidates Ebola virus origin and transmission during the 2014 outbreak. Science 2014, 345:1369–1372.

22. Park DJ, Dudas G, Wohl S, Goba A, Whitmer SL, Andersen KG, Sealfon RS, Ladner JT, Kugelman JR, Matranga CB, et al: Ebola Virus Epidemiology, Transmission, and Evolution during Seven Months in Sierra Leone. Cell 2015, 161:1516–1526.

23. Pruitt KD, Brown GR, Hiatt SM, Thibaud-Nissen F, Astashyn A, Ermolaeva O, Farrell CM, Hart J, Landrum MJ, McGarvey KM, et al: RefSeq: an update on mammalian reference sequences. Nucleic Acids Res 2014, 42:D756–763.

24. Luke S, Cioffi-Revilla C, Panait L, Sullivan K, Balan G: MASON: A multiagent simulation environment. Simulation-Transactions of the Society for Modeling and Simulation International 2005, 81:517–527.

25. Lindgreen S, Adair KL, Gardner P: An evaluation of the accuracy and speed of metagenome analysis tools. bioRxiv 2015.

26. Lauck M, Bailey AL, Andersen KG, Goldberg TL, Sabeti PC, O'Connor DH: GB virus C coinfections in west African Ebola patients. J Virol 2015, 89:2425–2429.

27. Mortazavi A, Williams BA, McCue K, Schaeffer L, Wold B: Mapping and quantifying mammalian transcriptomes by RNA-Seq. Nat Methods 2008, 5:621–628.

28. Jain M, Fiddes IT, Miga KH, Olsen HE, Paten B, Akeson M: Improved data analysis for the MinION nanopore sequencer. Nat Methods 2015, 12:351–356.

